# Little Evidence for Associations Between the Big Five Personality Traits and Variability in Brain Gray or White Matter

**DOI:** 10.1101/658567

**Authors:** Reut Avinun, Salomon Israel, Annchen R. Knodt, Ahmad R. Hariri

## Abstract

Attempts to link the Big Five personality traits of Openness-to-Experience, Conscientiousness, Extraversion, Agreeableness, and Neuroticism with variability in trait-like features of brain structure have produced inconsistent results. Small sample sizes and heterogeneous methodology have been suspected in driving these inconsistencies. Here, we tested for associations between the Big Five personality traits and multiple measures of brain structure using data from 1,107 university students (636 women, mean age 19.69±1.24 years) representing the largest attempt to date. In addition to replication analyses based on a prior study, we conducted exploratory whole-brain analyses. Four supplementary analyses were also conducted to examine 1) possible associations with lower-order facets of personality; 2) modulatory effects of sex; 3) effect of controlling for non-target personality traits; and 4) parcellation scheme effects. The analyses failed to identify any significant associations between the Big Five personality traits and variability in measures of cortical thickness, surface area, subcortical volume, or white matter microstructural integrity, except for an association between greater surface area of the superior temporal gyrus and lower scores on conscientiousness that explained 0.44% of the morphometric measure’s variance. Notably however, the latter association is largely not supported by previous studies. The supplementary analyses mirrored these largely null findings, suggesting they were not substantively biased by our choice of analytic model. Collectively, these results indicate that if there are direct associations between the Big Five personality traits and variability in brain structure, they are of likely very small effect sizes and will require very large samples for reliable detection.

## Introduction

Studies regarding the basic structure of individual differences in personality traits have yielded a relatively consistent five factor model, comprised of the higher-order dimensions of neuroticism, extraversion, agreeableness, conscientiousness, and openness-to-experience - each capturing a wide array of feelings, thoughts, and behaviors (Digman, 1990). Individuals high in neuroticism tend to perceive the world as distressing or threatening and frequently tend to experience negative emotions such as anger and anxiety. Extraversion reflects a tendency to be outgoing and assertive, to experience frequent positive moods, and to approach and explore one’s environment. Agreeableness reflects a tendency to be trusting and compassionate, and to prefer cooperation over conflict. Individuals high in conscientiousness tend to be organized and planful, and to follow socially prescribed norms of behavior. Individuals high in openness-to-experience tend to be curious and reflective, show an appreciation for art and culture, and tend to be very imaginative. These five traits can be further partitioned to a set of hierarchically lower-order facets, reflecting narrower, yet intercorrelated, sub-components of each broad dimension. In contrast to the consistency of the five factor model, studies of how the Big Five personality traits relate to underlying trait-like features of the brain have yet to identify consistent patterns despite a growing number of attempts (e.g., Bjørnebekk et al., 2013; Coutinho et al., 2013; DeYoung et al., 2010; Ferschmann et al., 2018; Hu et al., 2011; Kapogiannis et al., 2013; Liu et al., 2013; Lu et al., 2014).

Unlike the studies from which the Big Five were derived, most neuroimaging studies of personality traits have relied on relatively small samples (N<100). Importantly, a recent study has suggested that even samples of 300 participants may be too small to reliably detect associations between psychological phenotypes and brain morphometry (Kharabian Masouleh et al., 2019). Indeeed, consistent and replicable links have yet to emerge (reviewed in Allen and DeYoung, 2017; Yarkoni, 2015). This lack of statistical power has been further compounded by varied methodological and analytic approaches across studies. Here, we tested for associations between the Big Five personality traits and multiple features of brain structure in the largest sample to date (N=1,107). Notably, other than its size, our sample also had the advantage of being relatively homogeneous in age (18-22 years), which may affect associations between personality traits and brain structure (Ferschmann et al., 2018).

Brain morphometry was assessed by measuring cortical thickness (CT), surface area (SA), subcortical volume, and white matter microstructural integrity. Based on the radial unit hypothesis (Rakic, 1988, 2009), SA is driven by the number of radial columns, while CT reflects the density of cells within a column. CT and SA exhibit different developmental trajectories (Wierenga et al., 2014) and are affected by distinct genetic factors (Panizzon et al., 2009). Consequently, we examined associations with CT and SA separately rather than the coarser measure of gray matter volume, which is the product of these two measures.

To measure white matter microstructural integrity, we used fractional anisotropy (FA), which measures the directional diffusivity of water, with values ranging from 0 (isotropic diffusion) to 1 (anisotropic or directional diffusion). FA values have been associated with fiber diameter and density, degree of myelination, and fiber tract coherence (Basser, 1995; Basser and Pierpaoli, 1996; Beaulieu, 2002). Higher values of FA reflect a more organized or regular fiber tract pattern.

Additionally, we conducted surface-based parcellation analyses rather than traditional vertex- or voxel-based analyses to maximize spatial resolution otherwise lost to smoothing across tissue types (i.e., CSF, gray matter, and white matter) and anatomical regions (Coalson et al., 2018; Glasser et al., 2016). A parcellation-based approach further restricts the number of tests conducted to anatomically defined regions, thereby minimizing the number of comparisons conducted. We also explicitly controlled for race/ethnicity, because previous research has found it to be linked with brain structure (e.g., Brickman et al., 2008; Pfefferbaum et al., 2016; Xie et al., 2015) and personality (Foldes et al., 2008). Lastly, we used regression analyses with robust standard errors that accommodate non-normality in data, because previous research has indicated that the CT and SA of some brain regions may not be normally distributed (Patel et al., 2018).

Using the above general strategy, we conducted three related sets of analyses. First, we attempted to replicate the personality associations with brain morphometry reported by Hyatt et al. (2019), which represents the largest previously published study of the structural brain correlates of personality based on data from 1,104 participants, mostly twins and siblings, from the Human Connectome Project (age range: 22-36). Second, we conducted whole-brain exploratory analyses to examine all possible associations between the Big Five personality traits and SA, CT, subcortical volume, and FA. Third, in the hope of further informing future research in personality neuroscience, and to address previous findings and the possibility of parcellation scheme effects, we also conducted four supplementary analyses: we examined whether a) lower-order facets of the Big Five personality traits better correspond with brain structure (Bjørnebekk et al., 2013); b) associations differ by sex (Nostro et al., 2016); c) associations can be detected when controlling for non-target traits (e.g., DeYoung et al., 2010; Liu et al., 2013; Riccelli et al., 2017); and; d) a different parcellation scheme of the cortex affects the findings.

## Methods

### Participants

1330 participants (762 women, mean age 19.70±1.25 years) successfully completed the Duke Neurogenetics Study (DNS), which assessed a range of behavioral and biological traits among young adult, university students. The DNS was approved by the Duke University School of Medicine Institutional Review Board, and all participants provided written informed consent prior to participation. All participants were free of the following study exclusions: 1) medical diagnoses of cancer, stroke, diabetes requiring insulin treatment, chronic kidney or liver disease, or lifetime history of psychotic symptoms; 2) use of psychotropic, glucocorticoid, or hypolipidemic medication; and 3) conditions affecting cerebral blood flow and metabolism (e.g., hypertension). Current and lifetime DSM-IV (the Diagnostic and Statistical Manual of Mental Disorders) Axis I or select Axis II disorders (antisocial personality disorder and borderline personality disorder), were assessed with the electronic Mini International Neuropsychiatric Interview (Lecrubier et al., 1997) and Structured Clinical Interview for the DSM-IV Axis II subtests (First et al., 1997), respectively. Importantly, neither current nor lifetime diagnosis were an exclusion criterion, as the DNS seeks to establish broad variability in multiple behavioral phenotypes related to psychopathology. However, no individuals, regardless of diagnosis, were taking any psychoactive medication during or at least 14 days prior to their participation.

The current analyses of gray matter (i.e., CT, SA, and subcortical volume) were conducted on a subset of 1107 participants (636 women, mean age 19.69±1.24 years) for whom there was T1-weighted structural imaging data available post quality control procedures (see below) as well as personality questionnaire and genetic race/ethnicity data. Amongst this subset, 224 participants had at least one DSM-IV diagnosis. Based on self-report, there were 499 non-Hispanic Caucasians, 125 African Americans, 294 Asians, 71 Latino/as, 2 Pacific Islanders, and 116 multiracial or other participants in this subset.

White matter microstructure analyses were conducted on a further subset of 778 participants (443 women, mean age 19.67±1.25 years) for whom there was diffusion weighted imaging data available post quality control procedures (see below) as well as personality questionnaire and genetic race/ethnicity data. Amongst this subset, 156 participants had at least one DSM-IV diagnosis. Based on self-report, there were 351 non-Hispanic Caucasians, 92 African Americans, 213 Asians, 47 Latino/as, 2 Pacific Islanders, and 73 multiracial or other participants in this subset.

### Race/Ethnicity

Because self-reported race and ethnicity are not always an accurate reflection of genetic ancestry, an analysis of identity by state of whole-genome SNPs was performed in PLINK (Purcell et al., 2007). The first four multidimensional scaling components were used as covariates to reduce possible confounding effects of race/ethnicity. The decision to use only the first four components was based on an examination of a scree plot of eigenvalue, which showed that the eigenvalues became very similar after the fourth component.

### Personality

The 240-item NEO personality inventory revised (NEO-PI-R; Costa and McCrae, 1995), was used to assess the Big Five personality dimensions and their underlying facets: 1) Neuroticism (based on the anxiety, angry hostility, depression, self-consciousness, impulsiveness, and vulnerability facets); 2) Agreeableness (based on the trust, straightforwardness, altruism, compliance, modesty, and tender-mindedness facets); 3) Conscientiousness (based on the competence, order, dutifulness, achievement striving, self-discipline and deliberation facets); 4) Extraversion (based on the warmth, gregariousness, assertiveness, activity, excitement-Seeking, and positive emotions facets); and 5) Openness-to-Experience (based on the fantasy, aesthetics, feelings, actions, ideas, and values facets). Each facet was a sum of 8 items, and each personality trait was a sum of the facet scores (with certain items reverse coded as indicated). Participants rated the 240 items on a scale ranging from (0) *strongly disagree* to (4) *strongly agree.* The 6 lower order facets for each personality trait were modeled in our supplementary analyses. Internal consistency of the personality traits was assessed by Cronbach’s alpha as fair to good, ranging between .70 to .85.

### MRI Data Acquisition

Each participant was scanned using one of two identical research-dedicated GE MR750 3T scanners stationed at the same facility, the Duke-UNC Brain Imaging and Analysis Center (891 participants on scanner 1 and 216 participants on scanner 2. Additional details on the scanners can be found elsewhere: https://www.biac.duke.edu/facilities/scanners.asp). Each identical scanner was equipped with high-power high-duty cycle 50-mT/m gradients at 200 T/m/s slew rate and an eight-channel head coil for parallel imaging at high bandwidth up to 1 MHz. T1-weighted images were obtained using a 3D Ax FSPGR BRAVO sequence with the following parameters: TR = 8.148 ms; TE = 3.22 ms; 162 axial slices; flip angle, 12°; FOV, 240 mm; matrix =256×256; slice thickness = 1 mm with no gap (voxel size 0.9375×0.9375×1 mm); and total scan time = 4 min and 13 s. Following an ASSET calibration scan, two 2-min 50-s diffusion imaging acquisitions were collected, providing full brain coverage with 2-mm isotropic resolution and 15 diffusion weighted directions (10-s repetition time, 84.9 ms echo time, b value 1,000 s/mm2, 240 mm field of view, 90° flip angle, 128×128 acquisition matrix, slice thickness=2 mm). A variable that indicated the scanner that was used for each participant was included in all analyses as a covariate.

### MRI Data Processing

To generate regional measures of brain morphometry, anatomical images for each subject were first skull-stripped using ANTs (Klein et al., 2009), then submitted to Freesurfer’s (Version 5.3) recon-all with the “-noskullstrip” option (Dale et al., 1999; Fischl et al., 1999), using an x86_64 linux cluster running Scientific Linux. Of the 1321 participants who completed the high-resolution T1-weighted imaging protocol, 11 were excluded for the presence of motion-related or external artifacts, 4 were excluded for incidental findings, and 1 was unable to be processed with FreeSurfer. Additionally, the gray and white matter boundaries determined by recon-all were visually inspected using FreeSurfer QA Tools (https://surfer.nmr.mgh.harvard.edu/fswiki/QATools). This revealed small to moderate errors in gray matter boundary detection in 51 individuals who were consequently excluded.

CT and SA for 31 regions in each hemisphere, as defined by the Desikan-Killiany-Tourville atlas (Klein and Tourville, 2012), a modified version of the Desikan-Killiany atlas (Desikan et al., 2006), which was used in the Hyatt et al. (2019) study, were extracted using Freesurfer. The updated version of the atlas is meant to make region definitions as unambiguous as possible and define boundaries best suited to FreeSurfer’s classifier algorithm. To ensure that our exploratory analyses were not contingent on a specific parcellation scheme, CT and SA for 74 regions per hemisphere, as defined by the Destrieux atlas (Destrieux et al., 2010), were also extracted using Freesurfer. Additionally, gray matter volumes from eight subcortical regions (Cerebellum Cortex, Thalamus, Caudate, Putamen, Pallidum, Hippocampus, Amygdala, and Accumbens area) were extracted with Freesurfer’s subcortical segmentation (“aseg”) pipeline (Fischl et al., 2002). Estimated Total Intracranial Volume (ICV), total gray matter volume, cerebral white matter volume, and left and right hemisphere mean CT were also extracted from the “aseg” pipeline, and average whole-brain CT was calculated based on the estimates for the left and right hemispheres.

Diffusion weighted images were processed according to the Diffusion Tensor Imaging (DTI) protocol developed by the Enhancing Neuro Imaging Genetics Through Meta-Analysis (ENIGMA) consortium (Jahanshad et al., 2013; or http://enigma.ini.usc.edu/protocols/dti-protocols/). In brief, raw diffusion-weighted images underwent eddy current correction and linear registration to the non-diffusion weighted image in order to correct for head motion. These images were skull-stripped and diffusion tensor models were fit at each voxel using FMRIB’s Diffusion Toolbox in FSL (FDT; http://fsl.fmrib.ox.ac.uk/fsl/fslwiki/FDT), and the resulting two fractional anisotropy (FA) maps were linearly registered to each other and then averaged. Average FA images from all subjects were non-linearly registered to the ENIGMA-DTI target FA map, a minimal deformation target calculated across a large number of individuals (Jahanshad et al., 2013). The images were then processed using the tract-based spatial statistics (TBSS) analytic method (Smith et al., 2006) modified to project individual FA values onto the ENIGMA-DTI skeleton. Following the extraction of the skeletonized white matter and projection of individual FA values, tract-wise regions of interest, derived from the Johns Hopkins University (JHU) white matter parcellation atlas (Mori et al., 2005), were transferred to extract the mean FA across the full skeleton and average whole-brain FA values for a total of 24 (partially overlapping) regions across the two scans. All FA measures from the right and left hemispheres were averaged. Additionally, volume-by-volume head motion was quantified by calculating the root mean square (RMS) displacement of the six motion parameters (three translation and three rotation components), determined during eddy current correction for each pair of consecutive diffusion-weighted brain volumes. The resulting volume-by-volume RMS deviation values were averaged across all images, yielding a summary statistic of head motion for each participant to add to the FA analyses as a covariate, as previously recommended for DTI analyses (Yendiki et al., 2014).

### Statistical Analyses

We first attempted to replicate the significant associations between personality and brain morphometry reported by Hyatt et al. (2019; Table 1) at p<.005 (i.e., the significance threshold used in their paper). We next proceeded to conduct exploratory parcellation-based analyses across the whole-brain (31 SA regions, 31 CT regions, 8 subcortical regions, 24 FA measures, and total gray matter volume, cerebral white matter volume, whole-brain average FA, and whole-brain average CT) for each of the Big Five personality traits (a total of 5*98=490 tests). Lastly, to assess the robustness of our findings, we conducted four supplementary analyses: 1) whole-brain parcellation-based analyses of the Big Five personality facets; 2) whole-brain parcellation-based analyses of the Big Five personality traits for men and women, separately; 3) whole-brain parcellation-based analyses of each Big Five personality trait while controlling for the other four traits; and 4) whole-brain parcellation-based analyses of each Big Five personality trait, while using a different parcellation scheme of the cortex, specifically, the Destrieux atlas (Destrieux et al., 2010).

**Table 1.** Significant personality and brain structure (SA and CT) associations reported by Hyatt et al. 2019.

|  | CT | SA | Subcortical volume |
| --- | --- | --- | --- |
| **Neuroticism** | Left caudal middle frontal gyrus (+) | Left cuneus (-) |  |
|  | Left pars orbitalis (+) | Left pars triangularis (-) |  |
|  | Left pars triangularis (+) | Left superior parietal lobule (-) |  |
|  | Left superior frontal gyrus (+) | Right supramarginal gyrus (-) |  |
|  |  | Superior frontal gyrus (-) |  |
| **Openness** | Left rostral middle frontal gyrus (-) | Left inferior temporal gyrus (+) | Left caudate (+) |
|  | Left superior parietal lobule (-) |  |  |
| **Agreeableness** | Left caudal middle frontal gyrus (-) |  |  |
| **Extraversion** |  | Right superior frontal gyrus (+) |  |

Note. +/- indicate the direction of the associations (i.e., positive or negative respectively)

Analyses were conducted in R version 3.5.1 (R Core Team, 2018), with the packages “broom” (Robinson and Hayes, 2018), “tidyr” (Wickham and Henry, 2018), “dplyr” (Wickham et al., 2019), “lmtest” (Zeileis and Hothorn, 2002), “readr” (Wickham et al., 2018), and “sandwich” (Zeileis, 2004). Linear regression analyses with robust standard errors were performed with brain measures as outcomes, personality measures as independent variables, and sex, age, scanner, and four ancestry-informative genetic principal components as covariates of no interest. Notably, for all analyses, except the Hyatt et al., (2019) replication analyses, all brain morphometry measures were averaged across the two hemispheres, as there is no strong evidence to support a lateralization effect of personality on brain structure. ICV, average CT, and average FA, were used as additional covariates for analyses of subcortical volume and surface area, CT, and FA, respectively. For the FA analyses, which can be particularly sensitive to motion, head motion was also included as a covariate. The Big Five personality traits were standardized (M=0, SD=1) in SPSS version 25 before analyses. Variance explained (i.e., R^2^) by the independent variable of interest, when it is last in the regression, was calculated in R with the package “relaimpo” (Grömping, 2006). The “false discovery rate” (FDR) adjustment (Benjamini and Hochberg, 1995) was applied to correct for multiple comparisons with the p.adjust function in R.

## Results

Descriptive statistics for the personality and brain morphometry variables are available in Supplementary Table 1.

### Replication of Hyatt et al. (2019)

As reported in Table 2, none of the 15 associations that were significant at p<.005 in Hyatt et al. (2019) were significant in our analyses, even without correcting for multiple comparisons (i.e., using an uncorrected p<.05 threshold). As Hyatt et al. did not control for race/ethnicity, we also ran analyses without the genetic principal components to test whether these could account for the different results. Again, none of the associations remained significant after correcting for multiple comparisons, but three associations were significant at an uncorrected p<.05, although not necessarily in the same direction as found in Hyatt et al.: a positive association between the right supramarginal gyrus SA and neuroticism (b=24.047, SD=11.43, p=.036, R^2^=0.23%; this association was negative in Hyatt et al.); a negative association between the left pars orbitalis CT and neuroticism (b=-.013, SD=.005, p=.021, R^2^=0.33%; this association was positive in Hyatt et al.,), and a positive association between the left superior frontal gyrus CT and neuroticism (b=.0072, SD=.003, p=.02, R^2^=0.21%; this association was also positive in Hyatt et al.). When comparing the analyses with and without controlling for race/ethnicity (Table 2), it is noticeable that race/ethnicity can affect the obtained results.

**Table 2.** Regression estimates and standard errors from our attempt to replicate the associations reported by Hyatt et al. (2019) in all participants and within the three largest racial/ethnic subgroups from the Duke Neurogenetics Study.

|  | All participants (N=1107) |  |  | Without controlling for ethnicity (N=1107) |  |  | Caucasians (n=559) |  |  | Asians (n=336) |  |  | Africans Americans (n=143) |  |  |
| --- | --- | --- | --- | --- | --- | --- | --- | --- | --- | --- | --- | --- | --- | --- | --- |
|  | b | SD | R<sup>2</sup> | b | SD | R<sup>2</sup> | b | SD | R<sup>2</sup> | b | SD | R<sup>2</sup> | b | SD | R<sup>2</sup> |
| Left cuneus (SA) on Neuroticism | -4.263 | 7.610 | 0.02% | -0.787 | 6.977 | 0.00% | 5.879 | 11.276 | 0.04% | -9.265 | 12.485 | 0.13% | -26.246 | 18.164 | 1.11% |
| Left pars triangularis (SA) on Neuroticism | 6.955 | 6.536 | 0.07% | 9.857 | 6.095 | 0.14% | 15.075 | 10.201 | 0.29% | -6.948 | 11.284 | 0.08% | 35.2212<sup>^</sup> | 17.922 | 2.06% |
| Left superior parietal lobule (SA) on Neuroticism | 12.858 | 13.331 | 0.05% | 14.002 | 12.589 | 0.06% | 1.494 | 19.656 | 0.00% | 3.814 | 24.208 | 0.01% | 55.496 | 33.263 | 1.02% |
| Right supramarginal gyrus (SA) on Neuroticism | 20.087 | 12.314 | 0.16% | 24.0470* | 11.431 | 0.23% | 12.800 | 18.403 | 0.05% | 21.931 | 19.676 | 0.24% | 13.398 | 40.240 | 0.08% |
| Superior frontal gyrus (SA) on Neuroticism | 6.484 | 16.093 | 0.00% | 5.956 | 14.653 | 0.00% | 13.210 | 22.515 | 0.02% | 9.396 | 25.668 | 0.01% | 23.617 | 47.999 | 0.06% |
| Left inferior temporal gyrus (SA) on Openness | 14.579 | 11.062 | 0.09% | 16.364 | 10.347 | 0.11% | 28.9129<sup>^</sup> | 15.301 | 0.38% | -9.463 | 20.097 | 0.04% | 57.3691<sup>^</sup> | 29.662 | 1.48% |
| Right superior frontal gyrus (SA) on Extraversion | 19.777 | 18.350 | 0.03% | 19.780 | 17.327 | 0.03% | 4.725 | 26.898 | 0.00% | -57.200 | 34.859 | 0.29% | 79.808 | 55.550 | 0.44% |
| Left caudate on Openness | -17.406 | 12.350 | 0.11% | -17.243 | 11.604 | 0.11% | -39.6542* | 16.342 | 0.60% | -2.500 | 25.816 | 0.00% | 27.527 | 44.681 | 0.21% |
| Left caudal middle frontal gyrus (CT) on Neuroticism | 0.003 | 0.004 | 0.03% | 0.004 | 0.004 | 0.07% | 0.007 | 0.006 | 0.17% | 0.006 | 0.008 | 0.13% | 0.001 | 0.015 | 0.00% |
| Left pars orbitalis (CT) on Neuroticism | -0.008 | 0.006 | 0.14% | -0.0126* | 0.005 | 0.33% | -0.0162<sup>^</sup> | 0.008 | 0.54% | -0.006 | 0.010 | 0.08% | -0.018 | 0.019 | 0.55% |
| Left pars triangularis (CT) on Neuroticism | -0.003 | 0.004 | 0.02% | -0.004 | 0.004 | 0.07% | -0.005 | 0.006 | 0.08% | 0.004 | 0.007 | 0.06% | -0.010 | 0.012 | 0.34% |
| Left superior frontal gyrus (CT) on Neuroticism | 0.005 | 0.003 | 0.11% | 0.0072* | 0.003 | 0.21% | 0.001 | 0.004 | 0.01% | 0.0164** | 0.005 | 1.22% | -0.004 | 0.009 | 0.06% |
| Left rostral middle frontal gyrus (CT) on Openness | -0.002 | 0.004 | 0.02% | -0.001 | 0.003 | 0.01% | 0.000 | 0.005 | 0.00% | 0.002 | 0.007 | 0.01% | -0.012 | 0.012 | 0.42% |
| Left superior parietal lobule (CT) on Openness | -0.001 | 0.003 | 0.01% | -0.002 | 0.003 | 0.02% | 0.000 | 0.004 | 0.00% | -0.001 | 0.006 | 0.00% | -0.011 | 0.008 | 0.79% |
| Left caudal middle frontal gyrus (CT) on Agreeableness | -0.002 | 0.004 | 0.01% | -0.003 | 0.004 | 0.03% | -0.005 | 0.005 | 0.11% | -0.002 | 0.008 | 0.01% | 0.011 | 0.014 | 0.31% |

*Note.* CT=cortical thickness; SA=surface area; All participants=entire sample, controlling for 4 genetic principal components. In all analyses, sex, age and either mean CT or intracranial volume were included as covariates. <sup>^</sup>uncorrected p<.06, \* uncorrected p<.05, \*\*uncorrected p<.01 and FDR corrected p<.05

As the race/ethnic composition of our sample differed from the race/ethnic composition of the Human Connectome Project (HCP) sample on which the analyses of Hyatt et al. were based (i.e., 44.6% vs. 74.8% of non-Hispanic Caucasians, 11.4% vs. 15.1% of African-Americans, and 27% vs. 5.7% of Asian-Americans, which also included Native Hawaiian, or other Pacific Islander in the HCP), we also separately present the results from our three largest ethnic subsamples, as determined based on self-reports and genetic ancestry components, when available. Here, the sample sizes are larger because individuals with missing genetic data were also included based on self-reported race/ethnicity: non-Hispanic Caucasians (n=559), Asians (n=336), and African-Americans (n=143). As shown in Table 2, there were differences in the regression estimates between the groups, further supporting our decision to control for race/ethnicity (e.g., in Asians the association between the left superior frontal gyrus CT and neuroticism was positive and significant at an FDR corrected p value<.05, but it was somewhat negative in African Americans. This association was also significant at an uncorrected p<.05 in the mixed race/ethnicity sample, when race/ethnicity was not included as a covariate).

### Exploratory whole-brain analyses

The top associations (i.e., uncorrected p<.005) are reported in Table 3 along with their R^2^. Of all the associations between the Big Five personality traits and brain morphometry (CT, SA, subcortical volume, or FA) only one remained significant after the FDR correction for multiple comparisons: the association between the SA of the superior temporal gyrus and conscientiousness (b=-33.91, SE=8.66, p=9.55e-05, FDR adjusted p=0.047, R^2^=0.44%). All associations and related variance explained (R^2^) are further presented in Supplementary Table 2.

**Table 3.** Top results (p<.005) from whole-brain exploratory analyses between the Big Five personality traits and structural brain measures in the Duke Neurogenetics Study.

|  | **b** | **SE** | **p value** | **FDR adjusted**<br>**p value** | **R<sup>2</sup>** |
| --- | --- | --- | --- | --- | --- |
| **Superior temporal gyrus (SA) on Conscientiousness** | -33.9125 | 8.659488 | 9.55E-05 | 0.047 | 0.44% |
| **Thalamus Proper on Conscientiousness** | -54.9215 | 15.69009 | 0.000483 | 0.12 | 0.49% |
| **Postcentral gyrus (CT) on Openness** | 0.007176 | 0.002243 | 0.001413 | 0.23 | 0.45% |
| **Cerebral peduncle on Neuroticism** | -0.00165 | 0.000546 | 0.002643 | 0.32 | 0.82% |
| **Transverse temporal gyrus (SA) on Conscientiousness** | -3.85793 | 1.32717 | 0.003724 | 0.36 | 0.44% |

### Supplementary analyses

Our supplementary analyses (i.e., testing personality facets instead of the Big Five personality traits, conducting sex-specific analyses, using non-target personality traits as covariates or using a different cortex parcellation scheme; reported in Supplementary Tables 3-7) revealed that these generally null findings were not biased by our choice of analytic model. Only one association remained significant after the FDR correction for multiple comparisons across all the tests conducted in the current study (N=5640) - the association between the SA of the superior temporal gyrus and the dutifulness facet of conscientiousness (b=-39.50, SE=8.79, p=7.76e-06, FDR adjusted p=0.044; R^2^=0.62%), such that higher dutifulness was associated with reduced superior temporal gyrus SA.

## Discussion

In the current study, with the largest sample to date, we failed to identify robust links between the Big Five personality traits and multiple, trait-like features of brain structure. Several supplementary analyses, in which we tested the facets of personality, conducted separate analyses for men and women, included the non-target personality traits as covariates, and used a different parcellation scheme, confirmed these primary null findings. There was one exception: an association between greater SA of the superior temporal gyrus, an area involved in language perception and production, and lower scores on conscientiousness, and, more specifically, dutifulness. However, Bjørnebekk et al. (2013) who reported an association with the caudal part of the superior temporal gyrus, did so for scores on different facets of conscientiousness (achievement striving and self-discipline), and this association was not replicated in other studies (Hyatt et al., 2019; Lewis et al., 2018; Nostro et al., 2016), although an association with CT in this area was found to correlate with conscientiousness in Lewis et al., (2018). Consequently, this singular association in our current analyses should be treated with caution. Generally, the supplementary analyses suggested that the largely null results of the primary analyses did not depend on specific analytical or methodological choices. This is further supported by a recent large study which applied a voxel-wise approach and also did not find robust associations between personality traits and brain morphometry (Kharabian Masouleh et al., 2019).

There are several possible reasons for the lack of replicable associations between personality and brain structure, including the current failure to replicate associations identified by Hyatt and colleagues (2019). With regard to this specific failure, it is possible that unaccounted-for effects of family structure in the data derived primarily from twins and siblings (Van Essen et al., 2013), biased their observed associations. Additionally, even though we used a similar atlas (original study: the Desikan-Killiany atlas; our study: the updated Desikan-Killiany-Tourville atlas), a similar personality measure (original study: the 60-item NEO-FFI; our study: the 240-item NEO-PI-R), and a similar scanner (original study: the 3T Siemens Skyra; our study: the 3T GE MR750), it is possible that these small differences in data collection affected our results. However, if such differences account for the lack of replicability, this raises questions regarding the robustness of the original findings. More generally, most of the correlations reported by Hyatt et al. were smaller than .1, and, similarly, in our study almost all the R^2^ were smaller than 1%. This suggests that effect sizes for associations between personality traits and brain structure are likely to be very small and will require very large sample sizes to be reliably detected.

Furthermore, differences between and within samples may also limit replicability in personality neuroscience. Age is known to affect brain structure, and indeed has been shown to moderate associations between brain structure and personality (Ferschmann et al., 2018). Sex differences may also be relevant as has been shown by Nostro et al., (2016) and in our supplementary results, where, although an interaction by sex was not formally tested, the pattern of association between men and women were inconsistent. Our results also suggest that accounting for race/ethnicity may be advised when testing for personality-brain structure associations. Indeed, previous research has shown differences in brain structure as a function of race/ethnicity (e.g., Brickman et al., 2008; Pfefferbaum et al., 2016; Xie et al., 2015). For example, a different brain atlas than the one constructed based on Caucasians may be needed to accurately identify variability in structure from a different racial/ethnic population (Tang et al., 2010). Thus, it may be insufficient to simply control for race/ethnicity in analytic models.

Although we used the largest sample to date, included 240 items to assess personality, and employed different methodologies to test for the associations between the Big Five personality traits and brain structure, our study does have several limitations. First, we did not exhaust all the possible ways to assess personality. For example, an alternative classification approach is represented by personality “types,” which defines categories of individuals based on similar configurations of interacting traits. A large analysis of personality types indicated that there are 4 personality types that can be clustered based on scores on the Big Five personality traits (Gerlach et al., 2018). Thus, for example, someone low on neuroticism may also have an average or a high conscientiousness score, which may correspond differently with brain structure. Future studies could focus on such “types” in examining personality-brain structure associations. Second, our acquisition protocol precluded the application of more anatomically precise parcellation schemes (e.g., Glasser et al., 2016). Third, our sample of volunteer students at a top university may not be representative of the general population. Lastly, we did not examine brain function. The brain correlates of personality may be more readily identified in functional measures, such as functional connectivity. However, as functional MRI studies are often characterized by small sample sizes, here as well caution will be needed in the interpretation of findings until replicable findings emerge in large samples.

Our largely null findings echo comments made by Yarkoni (2015): “There is no guarantee that any particular psychometric model of individual differences in personality will map onto underlying biological process models in any straightforward way. In fact…a clear-cut relationship between the two is likely to be the exception rather than the rule.” As well as those by Kharabian Masouleh et al., (2019) that associations between psychological measures (including personality) and specific brain structures in a healthy sample are “highly unlikely” (Kharabian Masouleh et al., 2019). That said, small effect sizes and possible moderating effects of sex, age, and race/ethnicity suggest the possibility that with ever larger and more homogeneous samples reliable links between personality and trait-like features of the brain may yet emerge. The field of personality neuroscience may benefit from following the lead of genome-wide association studies that have, after many failed attempts with candidate gene studies and small samples (e.g., Avinun et al., 2018; Bosker et al., 2011), begun to reveal the genetic architecture of complex traits through massive samples (Plomin and von Stumm, 2018). The growth of shared imaging data through research consortia (e.g., the “enhancing neuroimaging genetics through meta-analysis” project [ENIGMA]) may allow for such gains in personality neuroscience sooner than later.

## ACKNOWLEDGEMENTS AND DISCLOSURES

We thank the Duke Neurogenetics Study participants and the staff of the Laboratory of NeuroGenetics. The Duke Neurogenetics Study received support from Duke University as well as US-National Institutes of Health grants R01DA033369 and R01DA031579. RA, ARK, and ARH received further support from US-National Institutes of Health grant R01AG049789. RA is also partly supported by a fellowship from the Jerusalem Brain Community. The Brain Imaging and Analysis Center was supported by the Office of the Director, National Institutes of Health under Award Number S10OD021480. The authors declare no competing financial or other interests.

## Notes

#### Summary of Updates

Added a Scanner covariate and variance explained to all analyses, and an additional supplementary analysis, testing personality associations with a different cortex parcellation scheme

## References

Allen, T.A., DeYoung, C.G., 2017. Personality neuroscience and the five factor model. In: Widiger, T.A. (Ed.), Oxford handbook of the five factor model. Oxford University Press, New York, pp. 319–352.

Avinun, R., Nevo, A., Knodt, A.R., Elliott, M.L., Hariri, A.R., 2018. Replication in Imaging Genetics: The Case of Threat-Related Amygdala Reactivity. Biological Psychiatry 84, 148–159.

Basser, P.J., 1995. Inferring microstructural features and the physiological state of tissues from diffusion-weighted images. NMR in Biomedicine 8, 333–344.

Basser, P.J., Pierpaoli, C., 1996. Microstructural and Physiological Features of Tissues Elucidated by Quantitative-Diffusion-Tensor MRI. Journal of Magnetic Resonance, Series B 111, 209–219.

Beaulieu, C., 2002. The basis of anisotropic water diffusion in the nervous system–a technical review. NMR in Biomedicine: An International Journal Devoted to the Development and Application of Magnetic Resonance In Vivo 15, 435–455.

Benjamini, Y., Hochberg, Y., 1995. Controlling the false discovery rate: a practical and powerful approach to multiple testing. Journal of the Royal statistical society: series B (Methodological) 57, 289–300.

Bjørnebekk, A., Fjell, A.M., Walhovd, K.B., Grydeland, H., Torgersen, S., Westlye, L.T., 2013. Neuronal correlates of the five factor model (FFM) of human personality: Multimodal imaging in a large healthy sample. Neuroimage 65, 194–208.

Bosker, F., Hartman, C., Nolte, I., Prins, B., Terpstra, P., Posthuma, D., Van Veen, T., Willemsen, G., DeRijk, R., De Geus, E., 2011. Poor replication of candidate genes for major depressive disorder using genome-wide association data. Molecular Psychiatry 16, 516.

Brickman, A.M., Schupf, N., Manly, J.J., Luchsinger, J.A., Andrews, H., Tang, M.X., Reitz, C., Small, S.A., Mayeux, R., DeCarli, C., 2008. Brain morphology in older African Americans, Caribbean Hispanics, and whites from northern Manhattan. Archives of Neurology 65, 1053–1061.

Coalson, T.S., Van Essen, D.C., Glasser, M.F., 2018. The impact of traditional neuroimaging methods on the spatial localization of cortical areas. Proceedings of the National Academy of Sciences 115, E6356–E6365.

Costa, P.T., McCrae, R.R., 1995. Domains and facets: Hierarchical personality assessment using the Revised NEO Personality Inventory. Journal of Personality Assessment 64, 21–50.

Coutinho, J.F., Sampaio, A., Ferreira, M., Soares, J.M., Gonçalves, O.F., 2013. Brain correlates of pro-social personality traits: a voxel-based morphometry study. Brain Imaging and Behavior 7, 293–299.

Dale, A.M., Fischl, B., Sereno, M.I., 1999. Cortical surface-based analysis: I. Segmentation and surface reconstruction. Neuroimage 9, 179–194.

Desikan, R.S., Ségonne, F., Fischl, B., Quinn, B.T., Dickerson, B.C., Blacker, D., Buckner, R.L., Dale, A.M., Maguire, R.P., Hyman, B.T., 2006. An automated labeling system for subdividing the human cerebral cortex on MRI scans into gyral based regions of interest. Neuroimage 31, 968–980.

Destrieux, C., Fischl, B., Dale, A., Halgren, E., 2010. Automatic parcellation of human cortical gyri and sulci using standard anatomical nomenclature. Neuroimage 53, 1–15.

DeYoung, C.G., Hirsh, J.B., Shane, M.S., Papademetris, X., Rajeevan, N., Gray, J.R., 2010. Testing predictions from personality neuroscience: Brain structure and the big five. Psychological Science 21, 820–828.

Digman, J.M., 1990. Personality structure: Emergence of the five-factor model. Annual Review of Psychology 41, 417–440.

Ferschmann, L., Fjell, A.M., Vollrath, M.E., Grydeland, H., Walhovd, K.B., Tamnes, C.K., 2018. Personality traits are associated with cortical development across adolescence: a longitudinal structural MRI study. Child Development 89, 811–822.

First, M.B., Gibbon, M., Spitzer, R.L., Williams, J.B.W., Benjamin, L.S., 1997. Structured Clinical Interview for DSM-IV Axis II Personality Disorders, (SCID-II). American Psychiatric Press, Washington, DC.

Fischl, B., Salat, D.H., Busa, E., Albert, M., Dieterich, M., Haselgrove, C., Van Der Kouwe, A., Killiany, R., Kennedy, D., Klaveness, S., 2002. Whole brain segmentation: automated labeling of neuroanatomical structures in the human brain. Neuron 33, 341–355.

Fischl, B., Sereno, M.I., Dale, A.M., 1999. Cortical surface-based analysis: II: inflation, flattening, and a surface-based coordinate system. Neuroimage 9, 195–207.

Foldes, H.J., Duehr, E.E., Ones, D.S., 2008. Group differences in personality: Meta-analyses comparing five US racial groups. Personnel Psychology 61, 579–616.

Gerlach, M., Farb, B., Revelle, W., Amaral, L.A.N., 2018. A robust data-driven approach identifies four personality types across four large data sets. Nature human behaviour 2, 735.

Glasser, M.F., Coalson, T.S., Robinson, E.C., Hacker, C.D., Harwell, J., Yacoub, E., Ugurbil, K., Andersson, J., Beckmann, C.F., Jenkinson, M., 2016. A multi-modal parcellation of human cerebral cortex. Nature 536, 171.

Grömping, U., 2006. Relative importance for linear regression in R: the package relaimpo. Journal of statistical software 17, 1–27.

Hu, X., Erb, M., Ackermann, H., Martin, J.A., Grodd, W., Reiterer, S.M., 2011. Voxel-based morphometry studies of personality: Issue of statistical model specification—effect of nuisance covariates. Neuroimage 54, 1994–2005.

Hyatt, C.S., Owens, M.M., Gray, J.C., Carter, N.T., MacKillop, J., Sweet, L.H., Miller, J.D., 2019. Personality traits share overlapping neuroanatomical correlates with internalizing and externalizing psychopathology. Journal of Abnormal Psychology 128, 1.

Jahanshad, N., Kochunov, P.V., Sprooten, E., Mandl, R.C., Nichols, T.E., Almasy, L., Blangero, J., Brouwer, R.M., Curran, J.E., de Zubicaray, G.I., 2013. Multi-site genetic analysis of diffusion images and voxelwise heritability analysis: A pilot project of the ENIGMA–DTI working group. Neuroimage 81, 455–469.

Kapogiannis, D., Sutin, A., Davatzikos, C., Costa, P.T., Resnick, S., 2013. The five factors of personality and regional cortical variability in the Baltimore longitudinal study of aging. Human Brain Mapping 34, 2829–2840.

Kharabian Masouleh, S., Eickhoff, S., Hoffstaedter, F., Genon, S., 2019. Empirical examination of the replicability of associations between brain structure and psychological variables. eLife.

Klein, A., Andersson, J., Ardekani, B.A., Ashburner, J., Avants, B., Chiang, M.-C., Christensen, G.E., Collins, D.L., Gee, J., Hellier, P., 2009. Evaluation of 14 nonlinear deformation algorithms applied to human brain MRI registration. Neuroimage 46, 786–802.

Klein, A., Tourville, J., 2012. 101 labeled brain images and a consistent human cortical labeling protocol. Frontiers in Neuroscience 6, 171.

Lecrubier, Y., Sheehan, D.V., Weiller, E., Amorim, P., Bonora, I., Sheehan, K.H., Janavs, J., Dunbar, G.C., 1997. The Mini International Neuropsychiatric Interview (MINI). A short diagnostic structured interview: reliability and validity according to the CIDI. European Psychiatry 12, 224–231.

Lewis, G.J., Dickie, D.A., Cox, S.R., Karama, S., Evans, A.C., Starr, J.M., Bastin, M.E., Wardlaw, J.M., Deary, I.J., 2018. Widespread associations between trait conscientiousness and thickness of brain cortical regions. Neuroimage 176, 22–28.

Liu, W.-Y., Weber, B., Reuter, M., Markett, S., Chu, W.-C., Montag, C., 2013. The Big Five of Personality and structural imaging revisited: a VBM–DARTEL study. Neuroreport 24, 375–380.

Lu, F., Huo, Y., Li, M., Chen, H., Liu, F., Wang, Y., Long, Z., Duan, X., Zhang, J., Zeng, L., 2014. Relationship between personality and gray matter volume in healthy young adults: a voxel-based morphometric study. PLoS ONE 9, e88763.

Mori, S., Wakana, S., Van Zijl, P.C., Nagae-Poetscher, L., 2005. MRI atlas of human white matter. Elsevier, Amsterdam.

Nostro, A.D., Müller, V.I., Reid, A.T., Eickhoff, S.B., 2016. Correlations between personality and brain structure: a crucial role of gender. Cerebral Cortex 27, 3698–3712.

Panizzon, M.S., Fennema-Notestine, C., Eyler, L.T., Jernigan, T.L., Prom-Wormley, E., Neale, M., Jacobson, K., Lyons, M.J., Grant, M.D., Franz, C.E., 2009. Distinct genetic influences on cortical surface area and cortical thickness. Cerebral Cortex 19, 2728–2735.

Patel, S., Patel, R., Park, M.T.M., Masellis, M., Knight, J., Chakravarty, M.M., 2018. Heritability estimates of cortical anatomy: the influence and reliability of different estimation strategies. Neuroimage 178, 78–91.

Pfefferbaum, A., Rohlfing, T., Pohl, K.M., Lane, B., Chu, W., Kwon, D., Nolan Nichols, B., Brown, S.A., Tapert, S.F., Cummins, K., 2016. Adolescent development of cortical and white matter structure in the NCANDA sample: role of sex, ethnicity, puberty, and alcohol drinking. Cerebral Cortex 26, 4101–4121.

Plomin, R., von Stumm, S., 2018. The new genetics of intelligence. Nature Reviews Genetics.

Purcell, S., Neale, B., Todd-Brown, K., Thomas, L., Ferreira, M.A., Bender, D., Maller, J., Sklar, P., De Bakker, P.I., Daly, M.J., 2007. PLINK: a tool set for whole-genome association and population-based linkage analyses. The American Journal of Human Genetics 81, 559–575.

R Core Team, 2018. R: A language and environment for statistical computing. R foundation for Statistical Computing, Vienna, Austria.

Rakic, P., 1988. Specification of cerebral cortical areas. Science 241, 170–176.

Rakic, P., 2009. Evolution of the neocortex: a perspective from developmental biology. Nature Reviews Neuroscience 10, 724.

Riccelli, R., Toschi, N., Nigro, S., Terracciano, A., Passamonti, L., 2017. Surface-based morphometry reveals the neuroanatomical basis of the five-factor model of personality. Social Cognitive and Affective Neuroscience 12, 671–684.

Robinson, D., Hayes, A., 2018. broom: Convert Statistical Analysis Objects into Tidy Tibbles. R package version 0.5.1.

Smith, S.M., Jenkinson, M., Johansen-Berg, H., Rueckert, D., Nichols, T.E., Mackay, C.E., Watkins, K.E., Ciccarelli, O., Cader, M.Z., Matthews, P.M., 2006. Tract-based spatial statistics: voxelwise analysis of multi-subject diffusion data. Neuroimage 31, 1487–1505.

Tang, Y., Hojatkashani, C., Dinov, I.D., Sun, B., Fan, L., Lin, X., Qi, H., Hua, X., Liu, S., Toga, A.W., 2010. The construction of a Chinese MRI brain atlas: a morphometric comparison study between Chinese and Caucasian cohorts. Neuroimage 51, 33–41.

Wickham, H., François, R., Henry, L., Müller, K., 2019. dplyr: A Grammar of Data Manipulation. R package version 0.8.0.1.

Wickham, H., Henry, L., 2018. tidyr: Easily Tidy Data with ‘spread()’ and ‘gather()’ Functions. R package version 0.8.2.

Wickham, H., Hester, J., Francois, R.F., 2018. readr: Read Rectangular Text Data. R package version 1.3.1.

Wierenga, L.M., Langen, M., Oranje, B., Durston, S., 2014. Unique developmental trajectories of cortical thickness and surface area. Neuroimage 87, 120–126.

Xie, W., Richards, J.E., Lei, D., Lee, K., Gong, Q., 2015. Comparison of the brain development trajectory between Chinese and US children and adolescents. Frontiers in systems neuroscience 8, 249.

Yarkoni, T., 2015. Neurobiological substrates of personality: A critical overview. In: Mikulincer, M.S., Cooper, P.R., Lynne, L.M., Randy, J. (Eds.), APA handbooks in psychology. APA handbook of personality and social psychology. American Psychological Association, Washington D.C., pp. 61–83.

Yendiki, A., Koldewyn, K., Kakunoori, S., Kanwisher, N., Fischl, B., 2014. Spurious group differences due to head motion in a diffusion MRI study. Neuroimage 88, 79–90.

Zeileis, A., 2004. Econometric computing with HC and HAC covariance matrix estimators.

Zeileis, A., Hothorn, T., 2002. Diagnostic checking in regression relationships. R news 2, 7–10.

